# A Common Methodological Phylogenomics Framework for intra-patient heteroplasmies to infer SARS-CoV-2 sublineages and tumor clones

**DOI:** 10.1101/2020.10.14.339986

**Authors:** Filippo Utro, Chaya Levovitz, Kahn Rhrissorrakrai, Laxmi Parida

## Abstract

We present a common methodological framework to infer the phylogenomics from genomic data, be it reads of SARS-CoV-2 of multiple COVID-19 patients or bulk DNAseq of the tumor of a cancer patient. The commonality is in the phylogenetic retrodiction based on the genomic reads in both scenarios. While there is evidence of heteroplasmy, i.e., multiple lineages of SARS-CoV-2 in the same COVID-19 patient; to date, there is no evidence of sublineages recombining within the same patient. The heterogeneity in a patient’s tumor is analogous to intra-patient heteroplasmy and the absence of recombination in the cells of tumor is a widely accepted assumption. Just as the different frequencies of the genomic variants in a tumor presupposes the existence of multiple tumor clones and provides a handle to computationally infer them, we postulate that so do the different variant frequencies in the viral reads, offering the means to infer the multiple co-infecting sublineages. We describe the Concerti computational framework for inferring phylogenies in each of the two scenarios. To demonstrate the accuracy of the method, we reproduce some known results in both scenarios. We also make some additional discoveries. We uncovered new potential parallel mutation in the evolution of the SARS-CoV-2 virus. In the context of cancer, we uncovered new clones harboring resistant mutations to therapy from clinically plausible phylogenetic tree in a patient.

## 1 Introduction

Deep sequencing genomic datasets contain intricate details that can be mined to reveal intra-patient heterogeneity present in disease states. The classic example that has been explored is the heterogeneity present in cancer, whether it be within a single tumor, across a patient’s metastatic sites, or a tumor’s evolution in response to treatment over the course of a disease. Interestingly, these same principles of heterogeneity can be explored in other scenarios that have similar sequencing data demonstrating different variant frequencies, including SARS-CoV-2 virus causing the COVID-19 infection. Evidence in several studies have highlighted the intra-host genomic diversity of SARS-CoV-2 [10, 11, 15, 23, 27]. As in cancer, the presence of different, heterogenic reads in a COVID-19 patient assumes the existence of multiple sublineages, or subclones, rather than the occurrence of recombination. Once these assumptions are established, the same tools and methodologies that are used to analyze tumor heterogeneity can be applied with a level of confidence to SARS-COV-2 datasets.

### Implications of viral heteroplasmy in COVID-19 patients

The novel SARS-Cov-2 coronavirus that appeared in the city of Wuhan, China, in late 2019 has caused a large scale COVID-19 pandemic, spreading to more than 70 countries. Broad sequencing efforts have been made in an effort to understand the natural evolution of this virus. Several studies published with SARS-CoV-2 sequencing data reveal different viral allele frequencies in the same patient. The most likely explanation for the presence of intra-patient heterogenic viral reads is the existence of different viral strains rather than recombination since the probability of a fully functional single stranded virus emerging after entering a cell and its subsequent disassembly and reassembly into a virion with a different sequence is low. Multiple viral strains infecting the same host has enormous clinical implications in terms of treatment, epidemiology, and the potential to overcome the pandemic and thus needs to be considered and analyzed. Variations in viral strains can harbor different resistance mechanisms, levels of transmissibility, response to therapy, and explain the large variation of symptomology. Even more important, treatment and vaccine success would rely on targeting the collection of strains present and not simply targeting one. It is for these reasons it is imperative that the research community consider the likely scenario that patients are coinfected with multiple strains.

### Implications of heterogeneity in tumors of cancer patients

The presence of multiple tumor clones in the same patient has significant treatment implications. Multiple mechanisms of resistance can exist in separate clones [22]. Drug targets can ‘disappear’ or develop over time [19, 26]. Alternate pathways can be inhibited by the introduction of new alterations [29]. Even gross phenotypes can change due to underlying genomic changes [1]. Thus, it is imperative that we continue to monitor patient tumor evolution over the course of a disease in order to optimize treatment protocols. Parallels of tumor evolution have been drawn to that of human evolution and thus similar tools and algorithms are being applied and adjusted to analyze cancer [21]. Phylogenetic trees are being constructed to capture the change occurring during the disease while subclonal structure is identified and analyzed for clinically relevant changes. Several algorithms have been proposed to capture tumor evolution using single cells [12, 25] however these tools do not account for tumor heterogeneity and thus do not pick up on all clones present in a given tumor. Most algorithms consider bulk tumor samples which has the advantage of integrating genetic information from many tumor cells but are challenged by the need to deconvolve the mixture of clones present in any given biopsy [4, 5, 7, 16, 18, 24]. Several of these methods have been adapted for multi-site sample integration but are not specific for longitudinal data [5, 9, 16, 18]. More recently there have been several tools developed that do integrate longitudinal (multi-time) sampling [14, 20]. Although these models are more accurate, they still are limited by their inability to deal with samples with large mutational burdens and are not designed for multi-site samples.

### Concerti overview

An informative analysis for SAR-CoV-2 would require a method to be able to perform fine-grain evaluations with the ability to differentiate between viral sequences. In addition, the method would need to be able to analyze longitudinal data to capture when co-occurrence transpires. In cancer, sequential liquid biopsies over the course of disease are becoming more common given the ease of collection, low cost, and greater ability to describe the complete disease profile vs. a subset of mutations present in distinct lesions. Therefore, it is imperative to establish tools that can manage/deconvolve mixed clonal samples, integrate multi-site and longitudinal sampling, and analyze large numbers of mutations with the same level of accuracy as low burden samples. We introduce Concerti a tool for inferring disease evolution phylogenies, at genomic scales, from multiple sites and multiple longitudinal DNA sequencing samples. One of the unique features of Concerti is that it generates *time-scaled trees*, i.e., it captures not only the birth and death of clones but also acquisition of alterations within the same clone over time. Concerti uses almost exclusively discrete optimization methods and has the flexibility to provide multiple possible solutions suggested by the patient data. To help with the interpretation of the results, the solutions are ordered by decreasing likelihood. Due to the absence of benchmark data, it is hard to perform a precise comparison of the different tools reported in literature. We provide a succinct summary of the capabilities of six classes of exemplar tools and highlight the elements of uniqueness in each approach. (Table 1).

**Table 1.** Exemplar phylogenetic methods for cancer and distinguishing features.

| Method | Input data | Data mode | Specific for multi-time data | Specific for multi-site data | Genomic scale | Time-scaled tree |
| --- | --- | --- | --- | --- | --- | --- |
| SCITE [12] | single cell | single value | ✗ | ✗ | N/A | ✗ |
| Pyclone [24] | bulk data | single value | ✗ | ✓ | ✗ | ✗ |
| CITUP [16] | bulk data | single value | ✗ | ✓ | ✗ | ✗ |
| Calder [20] | bulk data | single value | ✓ | ✗ | ✗ | ✓ |
| PhylogicNDT [14] | bulk data | distribution | ✓ | ✓ | ✓ | ✗ |
| Concerti | bulk data | distribution | ✓ | ✓ | ✓ | ✓ |

We demonstrate the accuracy of Concerti by reproducing and expanding on known results from literature. In particular, comparing with the results reported in [23] for the viral evolution model and expanding on them by discovering new homoplasies, while for the tumor evolution model using whole-exome sequencing data from patients [13, 22], Concerti constructs phylogenetic trees that accurately describe a tumor’s evolution while simultaneously highlighting new post-treatment subclones that likely confer resistance and may serve as new potential drug targets.

## 2 Method

### Viral Evolution Model

For computational purposes, we assume that all the virions of the same lineage have the same set of alterations (with respect to a reference). Since there is evidence of intra-patient variations with a wide range of allele frequencies [23, 28], we postulate that there is heteroplasmy due to possibly multiple sublineages evolving in this micro-environment. Since the coronavirus is a non-segmented positive RNA virus, we further postulate that it is very unlikely that any recombination occurs during the virus’s life cycle: attachment and entry, replicase protein expression, replication and transcription, assembly and release [6].

### Tumor Evolution Model

We assume the following model: tumors arise from an altered cell, accumulating additional alterations over time. These changes give rise to populations of cells termed in literature as *clones*. For computational purposes, we assume that all the cells in a clone have the same set of alterations. Furthermore, these clones may alter further over time. Thus multiple clones co-exist in a tumor and some may have an evolutionary advantage over the others within the tumor environment, allowing for growth or shrinkage of a clone over time.

### Terminology

The absence of recombination and the accumulation of variants over time are the two salient factors that facilitate a common methodology for both scenarios. We address the problem of inferring the evolution be it the coronavirus from multiple patients or tumor within a cancer patient. Furthermore for a cancer patient, the inferencing may be based on single or multiple DNA sequencing samples: the latter can be *multi-time*, i.e., at multiple timepoints, or can be *multi-site*, i.e., from different lesions possibly collected at the same time. We simply use the term *data point* for multi-patient (COVID-19) and multi-time, multi-site (cancer). The term *alteration* is applicable to any genetic event including, but not limited to, mutations, SNVs, etc. In this manuscript CCF (Cancer Cell Fraction) denotes the fraction of cancer cells bearing a mutation in a cancer sample [2]. For the purposes of our algorithm, CCF and VAF (Variant Allele Frequency) are indistinguishable and the precise method of determining alteration frequencies is outside the scope of this paper. For clarity of exposition we use VAF, in place of CCF or VAF, and SNV, in place of general alteration.

Furthermore, in the context of cancer, it is important to note the distinction between *clones* and *pseudoclones*. Clone is a biological entity and can be described as a population of indistinguishable cells. For our purposes, the nuclear DNA is identical for the population and thus a clone can be defined by a set of SNV’s (variant with respect to a reference or normal sequence). A pseudoclone on the other hand is a subset of these SNV’s. In practice they are a maximal collection of SNVs with identical (or similar) VAF values [18, 20, 24]: this value is termed *prevalence* in this paper. This collection of SNVs is meaningful under the assumption that identical VAF values implies these SNVs co-occur in a cell. Note that the converse is not necessarily true, i.e., multiple SNVs within a cell may have varied VAF values. Thus pseudoclone is an algorithmic artefact while a clone is simply the union of some finite pseudoclones. For example in Fig. 4, the distinct colors denote the different pseudoclones, but the biological clone is the union of all the subclones in the path to the root of the evolutionary tree. Hence the (biological) brown clone in the subcutaneous soft tissue, for instance, is actually the the union of the SNVs that define the brown pseudoclone, the orange pseudoclone, the cyan pseudoclone and the green pseudoclone. Hence, for mathematical preciseness, we use the term pseudoclones in the Method sections. To avoid clutter, we use the term clone, in place of pseudoclone, in the Result Section.

With a slight abuse of terminologies, a pseudoclone corresponds to a sublineage in the context of virions and a clone corresponds to a lineage. To avoid clutter, we use the terms sublineage and lineage interchangeably.

#### 2.1 Method Assumptions

We make the following assumptions.

##### Assumption 1. [Infinite Sites Model]

A majority of the alterations satisfy the following:

1. irreversible, i.e., once the alteration occurs the reverse of turning it back to its original state does not occur (no back mutation).
2. unique, i.e., the same alteration does not occur elsewhere in the tumor (no parallel mutation).

The topology of the evolution is a tree. The Occam’s Razor Principle suggests the perfect phylogeny assumptions used most commonly in literature [3]. However, it is important to note that some exceptions to this property of alterations may occur in practice. This is handled as an exception in our algorithm. These exceptions indicate parallel mutation. This is also referred to as *homplasy*.

##### Assumption 2. [Alteration distribution]

Most of the alterations follow i.i.d. (uniform) distribution.

Again, for algorithmic purposes, it is reasonable to assume that tumor clones would follow the same principles as the individual alteration. But a clone, unlike an alteration, may die, i.e., may be selected against and overrun by other clones. So a clone may change in composition over time, i.e. more alterations can be added to the clone (but, not removed due to Assumption 1) for a majority of the alterations. Different selection pressures are in effect on the different clones, whose effect is manifested in the size of the clones: the clone can either grow or shrink in size reflected as an increasing or decreasing CCF value respectively. Thus the following:

##### Assumption 3. [Tumor Clone dynamics]

Over time, a clone may

1. change in composition / size (additional alterations but not lose alterations)
2. change in prevalence values (increase or decrease)
3. die or a new clone may be born.

#### 2.2 Method Overview

Concerti takes as input the VAF or CCF distribution for each alterations in all *n* data points which can be multi-patient (COVID-19) or multi-time or multi-site data of a patint (cancer).

See Fig. 1 for an overview of Concerti. Based on our assumptions, the method has two major phases.

**Figure 1.**
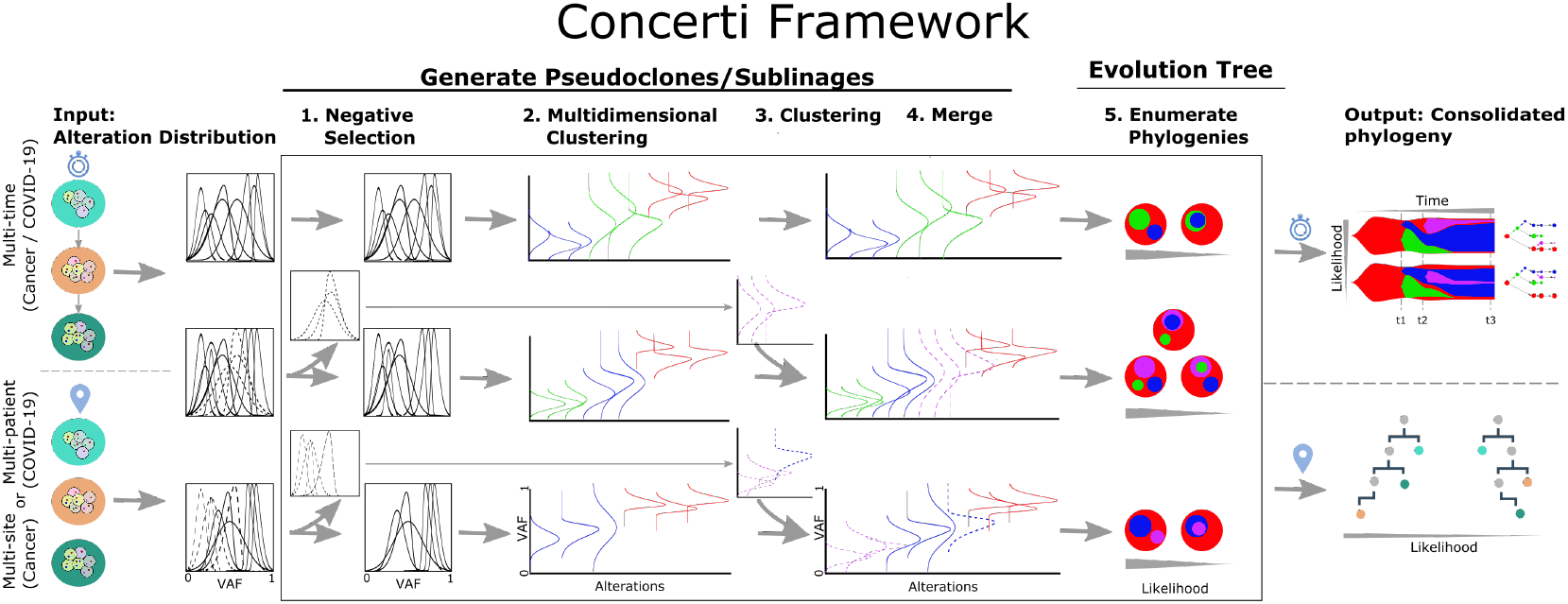
Schematic of the Concerti Framework. Given a set of multi-patient (COVID-19) or multi-site, multi-time (cancer) genomic samples, the algorithm analyzes the underlying alteration frequency distribution as input and performs a (1) negative selection to filter *appearing* alterations. A (2) multidimensional clustering is done to identify pseudoclones/lineages that will then be enriched by a (3) single sample clustering that (4) merges alterations that were initially negatively selected. (5) All potential phylogenies are generated and assessed for compatibility. Finally the set of consolidated phylogenetic structures over time or site are output with likelihood scores.

1. **Phase I**. We first identify the pseudoclones/sublineages across all the data-points. However, the pseudoclones are not identical due to the clonal dynamics (Assumption 3).

a. Due to Assumption 3(3), we gate the alterations that are present in all the samples. This results in some alteration being filtered out and the step is called Negative Selection in the outline. We cluster the filtered alterations separately for each sample.
b. The pseudoclones are preserved in the samples, albeit with some dynamics (Assumption 3(2)). We carry out a multi-dimensional clustering, across all samples, based on the values or distributions of the alterations.
c. We appropriately merge the clusters of the above two steps to obtain the pseudo-clones/sublineages.
2. **Phase II**. This has two steps: in the first we deduce the phylogenies of each sample separately and in the second we relate them with each other.

a. The sizes of the pseudoclones/sublineages in each data-point admits possibly multiple phylogenies. We enumerate the admissible phylogenies associating a probability with each based on Assumption 2.
b. Next we consolidate the trees from the multiple data points. This captures the topology as well as the clonal dynamics. Concerti offers two types of visualizations: one that effectively captures the change-in-composition dynamics of the clones (as a tree) and the other that captures the change-in-size and birth/death of clones effectively (fishplots [17]). The multiple possible solutions suggested by the data is output in decreasing order of probabilities, making it easier for interpreting the results.

##### Exception Handling

Real data is sometimes notoriously perplexing, either due to errors in sample or CCF distribution estimation or simply the infraction of some of the assumptions enumerated in the last section. In practice, in refractory cases we handle the exceptions by relaxing the minimum of assumptions by consulting with the domain experts.

#### 2.3 Phase I: Generate Pseudoclones/Sublineages

We define the distance between a pair of CCF distribution *g*_1_ and *g*_2_ as

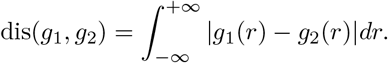

However, for algorithmic efficiency, we use the similarity measure defined as

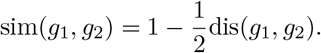

Thus identical distributions have a similarity of 1 while distinct distributions have similarity value zero. In practice, since the probability density function is specified as a discrete set of pairs of values and its probability, we compute the similarity as follows:

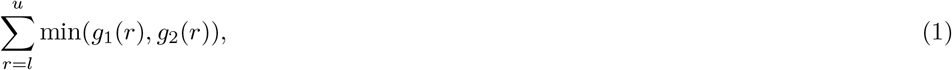

where [0 <= *l*, *u* <= 1.0] is the maximal interval where both *g*_1_ and *g*_2_ have non-zero values. We first perform a negative selection where alterations not present in all samples are removed. Let this set of filtered alterations be *S*_*a*_. Let the remaining set of alterations be *S*_*b*_. We carry out a hierarchical clustering [8] of the alteration set *S*_*b*_ using the similarity function (1). We cluster *S*_*a*_ for each sample separately and then merge the resulting multi-dimensional clustering of *S*_*b*_ with the sample-by-sample clustering of *S*_*a*_ to produce the pseudoclones. The *prevalence* of a pseudoclone is approximated by the mean of the mean value of each constituent alteration (CCF) distribution.

#### 2.4 Phase II: Generate Evolution Tree(s) of Pseudoclones/Sublineages

##### 2.4.0.1 Enumerate Admissible Trees

The question of how multiple sets of alterations (pseudoclones/sublineages) at a data point stack up against each other is almost completely left to the computational methods.

Fig. 2 shows a simple example. Assume without loss of generality (WLOG) u=1.0. Prevalence values 0 ≤ *a*, ≤ *b* 1 can be viewed as some sub-intervals (*sticks*) of [0, 1] of lengths *a* and *b* respectively. Then in a cell-population realization either sticks *A* (with prevalence value *a*) and *B* (with prevalence value *b*) are nested or disjoint but may not straddle. To remove possible computational artefacts, pseudoclones with prevalence *v* < 0.05 and less than 3 SNVs for all samples are discarded. Finally, given a pseudoclone A with a prevalence value *a*, then for each *x* pseudoclone nested directly in A with prevalence *v*_*x*_, the sum of their prevalence is Σ_*x*_ *v*_*x*_ ≤ *a*. With bulk-sequencing (i.e. no viral isolates or single-cell), the multiple possible scenarios cannot really be teased apart. But using Assumption 2, the probability of each admissible tree can be estimated.

**Figure 2.**
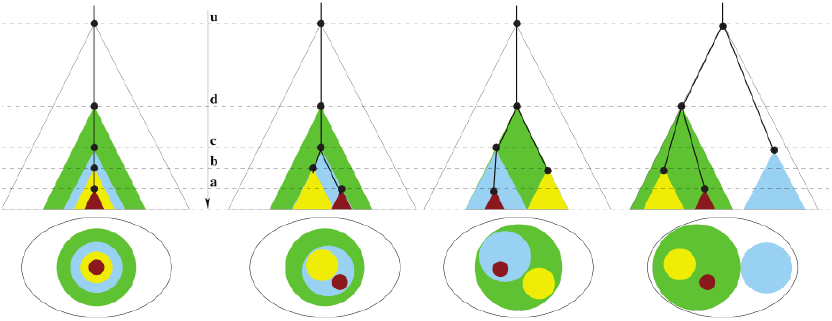
Let U (white), D (green), C (cyan), B (yellow), A (brown) be pseudoclones/sublineages with prevalence values: 1.0 *u* > *d* > *c* > *b* > *a* > 0 respectively. The top row shows 4 possible evolution trees where the time axis is the *molecular clock*. The bottom row shows the single time-point “fishplot” as appropriately stacked disks. Notice that the leftmost phylogeny suggests that there exist some cells/virion with both A and B alterations while all the other three suggest that there exists no such cell/virion.

We present a recursive algorithm, called Stick-Stack (Algorithm 1), that enumerates all possible ways of stacking the sticks (or sub-intervals) corresponding to pseudoclones/sub-lineage. The algorithm computes the probabilities of the trees using Assumption 2. In the output of Stick-Stack (*A, B*) denotes nesting of *B* in *A*. Stick-Stack call is initiated as (1, *ϕ*, *pr* = 1.0, *v*_1_ >…> *v*_*k*_) with the *n* prevalence values of the *k* pseudoclones/sublineages and the output is a set of trees where each tree *tr* is a collection of (parent,child) pairs, with probability *pr*. It is easy to verify from the algorithm description below that the probabilities of all the admissible trees sum up to 1.

##### 2.4.0.2 Mutual Comparison

While a single data point may suggest the relative relationship between pseudoclones, clonal dynamics can only be captured from multiple data points, be it multi-time or multi-site.

**Algorithm 1:**
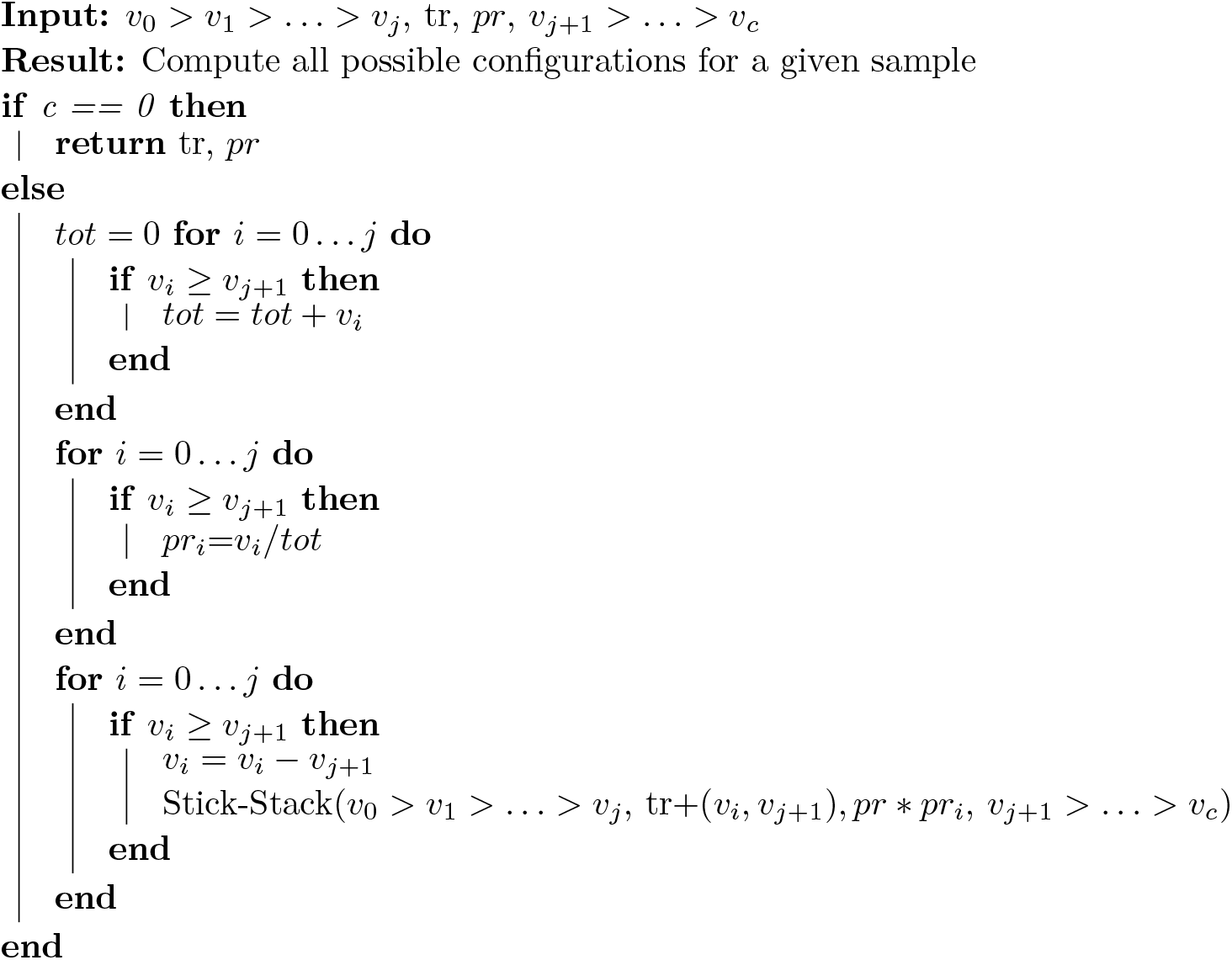
Stick-Stack: Given the prevalences, the algorithm enumerates all possible admissible trees, each with an estimated probability of occurrence.

###### Observation 1. [Clone Dynamics]

Based on Assumption 3 the clusters of filtered alterations of Step 1 of Phase I provide the clone dynamics.

a. If such a cluster merges with the multidimensional cluster, then this indicates a change in composition of the pseudoclone and provides labels for the edges of the evolution phylogeny.
b. If a new cluster is generated (i.e., it does not merge with clusters from the multi-dimensional clustering) then this indicates the birth of a new pseudoclone.

A clone acquiring new alterations (case a. above) is shown as asterisk in the tumor phylogeny in Fig. 5 and 6. The birth and death of clones (case b. above) is also illustrated in the same figure.

The mutual comparisons reveal whether the different samples are related or independent. When related, it is possible to reconstruct *consolidated* tree(s) that capture the evolution across the the multiple data points. Stick-Stack algorithm produces each tree as a set of two-tuples corresponding to each edge as (parent,child), where the parent and child are both pseudoclones. Formally, if there exist *k* > 0 parent-child pairs as (*C*_0_, *C*_1_), (*C*_1_, *C*_2_), …, (*C*_*k−*1_, *C*_*k*_), then *C*_0_ *precedes C*_*k*_ or *C*_0_ ≺ *C*_*k*_.

###### Definition 1. [Incompatible]

Let 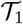 and 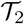 be two trees with three sets of (possibly empty) pseudoclones: *A*_*i*_ that occur in both 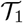 and 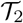; *D*_*i*_ that occur in 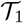, but not in 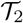, and, *B*_*i*_ that occur in 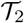 but not in 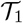.

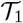 and 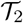 are incompatible if at least one of the following conditions does not hold:

1. WLOG if *A*_1_ ≺ *A*_2_ in 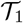 then 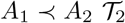.
2. WLOG each *D*_*i*_ is of the type (−, *D*_*i*_) and if *D*_*i*_ has a child then it occurs as (*D*_*i*_, *D*_*j*_) in 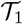.
3. WLOG each *B*_*i*_ is of the type (−, *B*_*i*_) and if *B*_*i*_ has a child then it occurs as (*B*_*i*_, *B*_*j*_) in 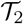. Once all possible configurations of the pseudoclone for any given time point are generated independently, the next step is to extract the possible compatible tree between all time points.

###### Definition 2. [Consolidated phylogeny]

Let 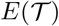 be the set of the two tuples (parent,child) of 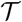. If 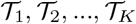 are mutually compatible, then **T**, the consolidated phylogeny of the *K* trees, is defined by the following set

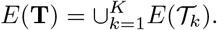

Notice that by conditions 2 and 3 of the incompatible definition above, **T** does not have any nodes with multiple parents and **T** is a tree.

Let 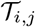 be the *j*th compatible tree at data point *i* with probability *p*_*i,j*_ as estimated by Stick-Stack. Then the the relative probability of a compatible evolution tree 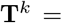 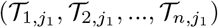 over the *n* data points is given by

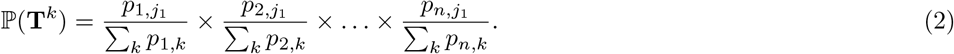

Note that Σ_*k*_ ℙ(**T**^*k*^) = 1 where *k* is over all possible compatible configurations. Thus the probability of a **T**^*k*^ may be underestimated. However, it preserves the ordering of the possible multiple solutions which is used here.

For a concrete example, consider Fig. 4. The four sites are labeled as subcutaneous soft tissue (subcu), brain, liver1 and liver2. For each site, Concerti produces exactly one tree, with eight pseudoclones across all the four sites: green (G), cyan (C), orange (O), purple (P), ash (A), yellow (Y), red (R), brown (B). Then Stick-Stack produces the following four trees:

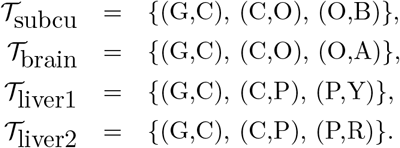

It can be verified that the four trees are mutually compatible. Then the unique consolidated phylogeny is given by

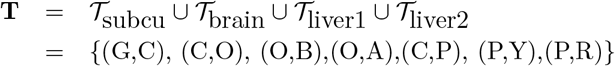

**T** has only one connected component suggesting that all the data points are genetically related.

In multi-time data, a consolidated tree **T** is stretched out with the given time points appropriately marking the tree (see Figs.5, 6 for example). Additionally, a fishplot is output to visualize the dynamics of the pseudoclones (growth or shrinkage, including birth and death).

## 3 Results

We applied Concerti on publicly available COVID-19 sequencing data as well as multi-site and multi-time cancer sequencing data (see Data Availability Section).

### 3.1 COVID-19 data

For our study we sought COVID-19 patient samples with access to the raw reads in order to assess the alterations at varying allele frequencies. Using an established reference MN908947.3 obtained from the ‘first’ patient sequenced in Wuhan, we found forty one distinct variants in twenty one patient samples (see the Data Availability Section for details on the patient samples and the variant calling pipeline). Eighteen of the patients were also analyzed in [23], albeit the variants therein were derived based on a different reference sequence and protocol. Hence, the set of variants do not exactly match the set we obtain. Our data set has three additional patients (who are sellers or deliverymen from the Wuhan seafood market) [30]. We applied Concerti to the data and the resulting phylogeny is shown in Fig. 3. Note that all samples with dark blue sublineage in the figure were collected in the USA, the one with the dark-grey sublineage (SRR11278092) was collected in Nepal, while the remaining were collected in China. The figure shows the other lineages that were identified. The phylogeny is not fully resolved based on the variants of this set of patients; this is shown as clusters of patients in two internal and one leaf node in the tree. The phylogeny also uncovers three parallel mutation events (shown as transversal dashed lines with the corresponding color): 404:A>T (raspberry), 29039:A>T (grey) and 4229:A>C (green). The first two were reported in [23] while the third is discovered in this study.

**Figure 3.**
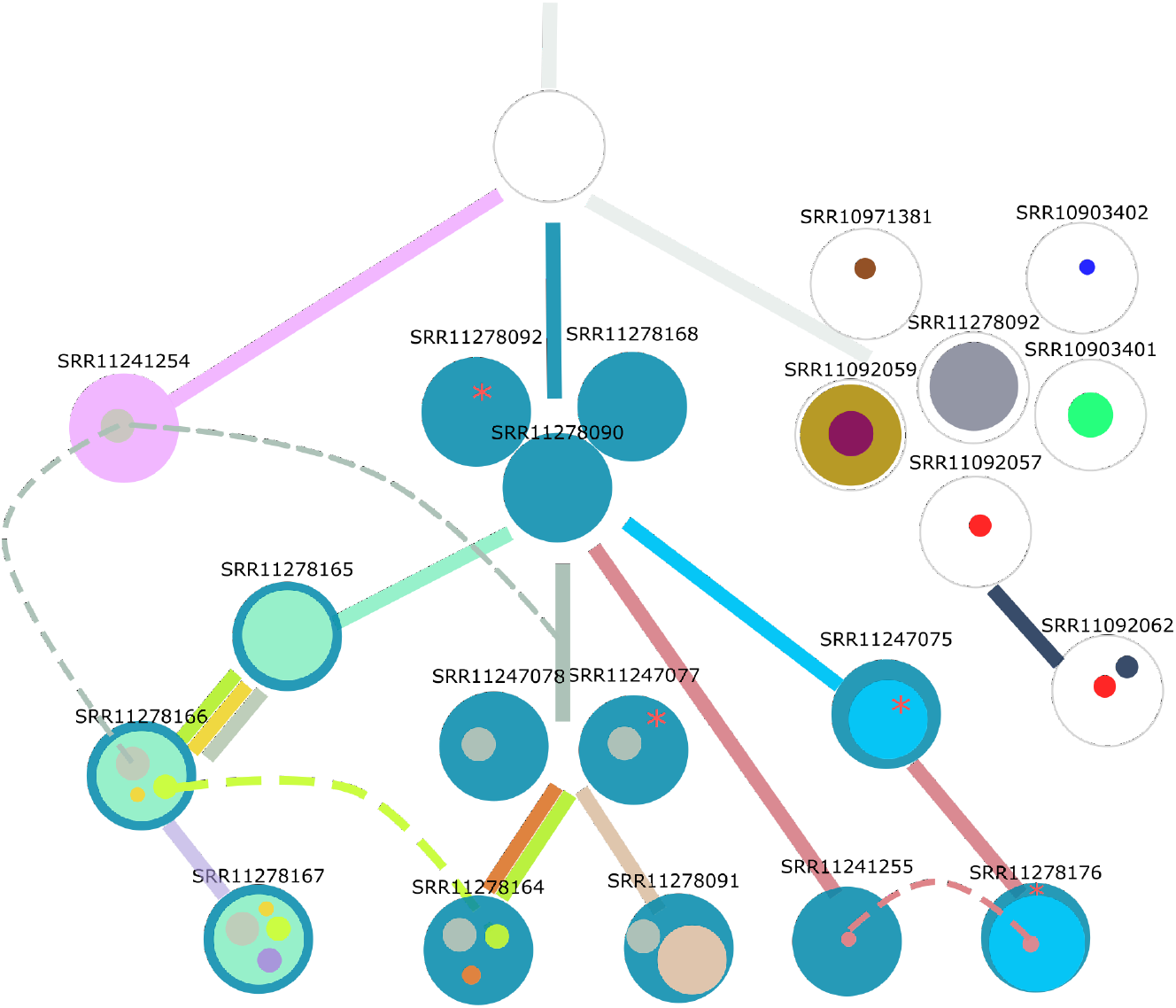
The 21 COVID-19 patients are shown at different internal and leaf nodes in the phylogeny as stacked disks of different colors. Each color corresponds to a distinct sublineage identified by a set of alterations and the size is roughly proportional to its observed prevalence value. Where possible, the edges of the phylogeny are colored by the emerging sublineage(s). When a node has multiple individuals, it indicates that there is not enough evidence to delineate the distinctions in the phylogeny. The three homoplasies (parallel mutations) are shown by dashed transversal lines. While in two (raspberry, green colors) the alteration event occurred at least twice, in the third (gray color) the alteration occurred at least three times. Furthermore, if the date of collection of a sample at a child node precedes the date at a parent node, it is within a window of a week.

### 3.2 Cancer data

Using Concerti, we analyzed three patients, one sampled across multiple sites (Fig. 4) and two sampled over time (Figs. 5 and 6). We first applied Concerti to a multi-site case, GI1, a 53 year old male with metastatic colon cancer who was part of a rapid autopsy study where multiple metastatic samples were taken at the time of death. Additional clinical details and the description of the sequencing method can be found in [22]. Samples were taken from different anatomical sites including lesions in the liver, brain, and subcutaneous soft tissue. Time of lesion development was not documented radiologically and thus no longitudinal time-ordering of the samples could be performed. Concerti’s generated tumor evolution tree gives a clinically plausible explanation as to the mutational development of this disease (Fig. 4). The phylogenetic tree characterizes several truncal clones shared across all samples (green and cyan) and then identifies two sibling clones (orange and purple) that are tissue specific. One clone captures both liver samples and contains the KRAS p.G12S allele. The other clone, which contains the ELF3 p.S229R allele, goes on to develop two daughter clones each specific to the brain or subcutaneous soft tissue. Thus, Concerti’s integration of multi-site samples enables the phylogenetic tree to capture a tumor’s broad spatial heterogeneity and allows for a treatment course to be designed to be locally or broadly targeted.

**Figure 4.**
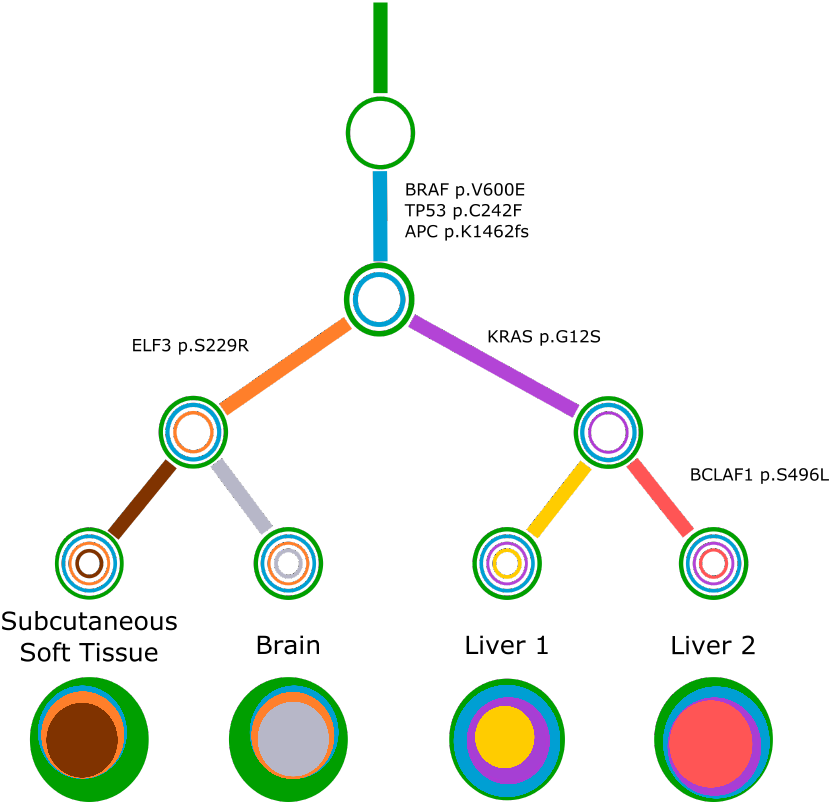
Concerti tumor evolution tree **T** for patient GI1. Tumor evolution tree **T** for colon cancer patient GI1 multi-site data. The edges of the **T** are labeled by the known cancer genes and the colors denote the distinct pseudoclones estimated by Concerti. Leaf nodes represent each of the distinct lesion sites. The single site trees 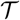 are shown at the bottom as stacked discs and the sizes are proportional to the prevalence values.

**Figure 5.**
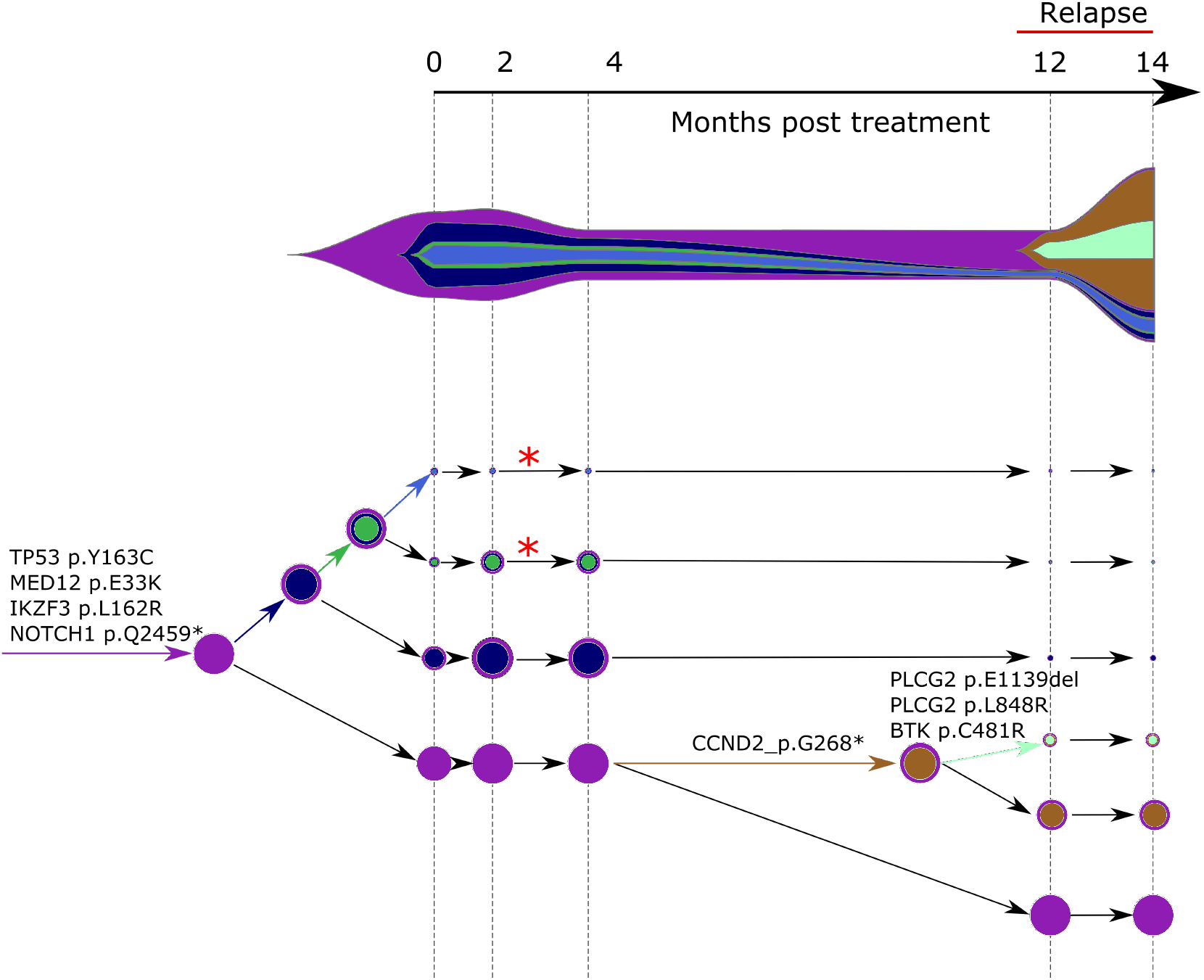
Concerti fishplot and tumor evolution tree **T** for patient CLL1 multi-time data. The fishplot width corresponds to approximate tumor size using ALC (absolute lymphocyte count) values. Clones are colored and sized proportionally to their prevalence. The corresponding tumor tree is aligned by timepoint and highlights to the birth of brown clone, which occurred prior to the annotation of clinical relapse. Node sizes correspond to prevalence. The edges of the **T** are labeled by the known cancer genes and the colors denote the distinct pseudoclones estimated by Concerti. Red asterisks indicates acquisition of new alterations to clone.

**Figure 6.**
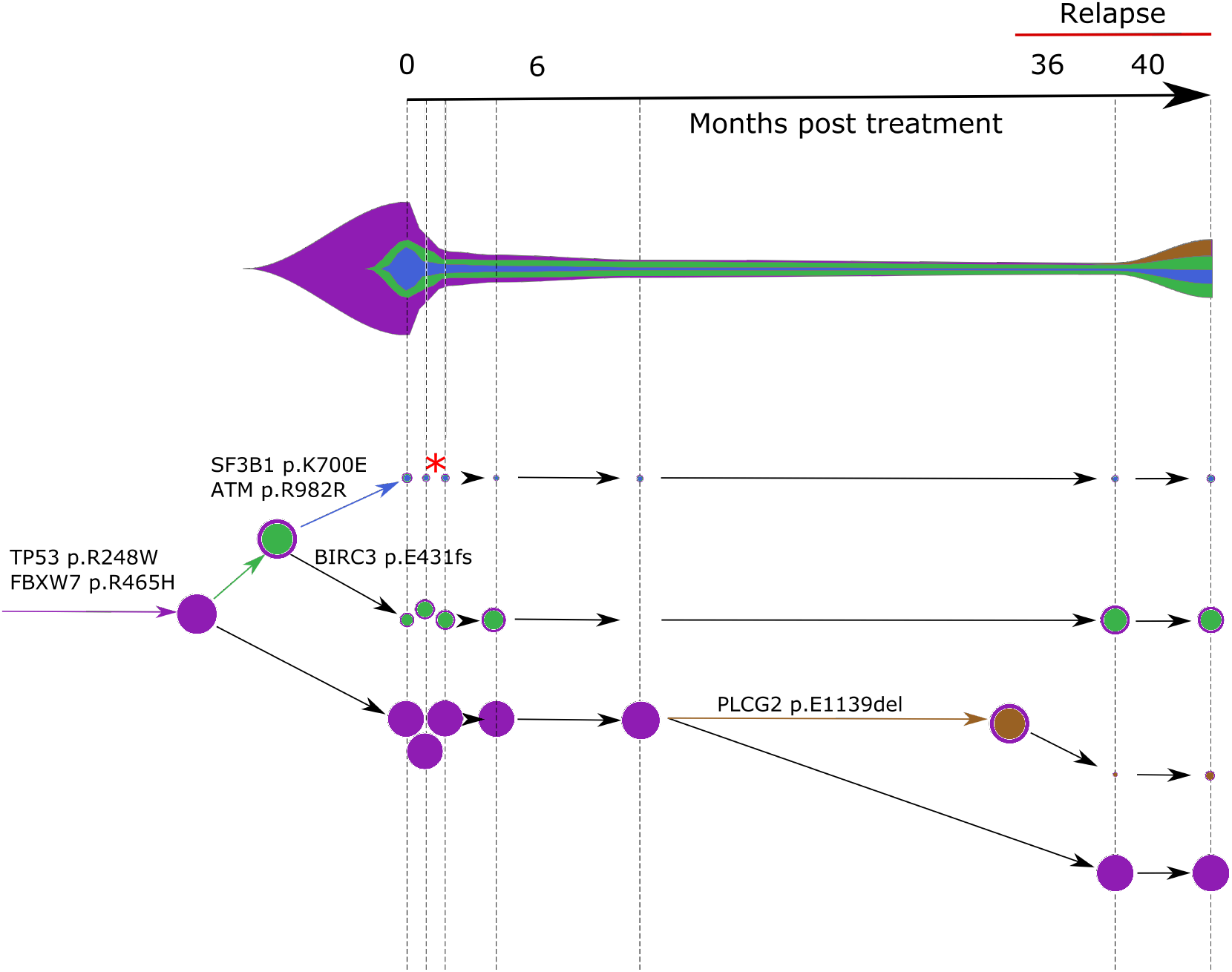
Concerti fishplot and tumor evolution tree **T** for patient CLL2 multi-time data. The fishplot width corresponds to approximate tumor size using ALC values. Clones are colored and sized proportionally to their prevalence. The corresponding tumor tree is aligned by timepoint and highlights to the birth of brown clone, which occurred prior to the annotation of clinical relapse. Node sizes correspond to prevalence. The edges of the **T** are labeled by the known cancer genes and the colors denote the distinct pseudoclones estimated by Concerti. Red asterisk indicates acquisition of new alterations to clone.

We also applied Concerti to longitudinal sequencing data from two relapsed chronic lymphocytic leukemia (CLL) patients. Patient CLL1 had five biopsies taken over the course of treatment with ibrutinib and rituximab and relapsed 12 months after treatment initiation (Fig. 5). Before treatment, the dominant clone contained mutations in several known cancer genes including TP53, MED12, IK2F3, and NOTCH1. Two small clones (red asterisks) continued to evolve as evidenced by the acquisition of additional mutations after two months post-treatment. The fishplot highlights the correspondence between the emergence of a resistant clones and the increase in tumor size. Concerti’s time-scaled phylogenetic tree and fishplot captures the birth of this clone, before relapse was clinically documented, that harbored three mutations in genes associated with resistance to ibrutinib, including BTK, PLCG2, and known cancer driver CCND2.

The second CLL patient Concerti analyzed was similarly treated with and developed resistance to ibrutinib. (Fig. 6). For patient CLL2, seven blood biopsies were taken over the treatment course including before-treatment, on-treatment, and at time of relapse. Several truncal mutations in known cancer genes were identified in the pre-treatment samples, including TP53 and FBXW7. After initiation with ibrutinib, a clone with a BIRC3 mutation (green) increased in prevalence. At the time of relapse, Concerti’s phylogenetic tree identifies the emergence of a new clone harboring a mutation in PLCG2, a known mechanism of resistance to ibrutinib therapy, and which goes on to grow in prevalence. Clones with ATM and SF3B1 did not have noticeable clonal dynamics during the treatment or relapse intervals suggesting they are not selected for under ibrutinib therapy. In both CLL patients, the birth of these resistant clones in response to treatment was only able to be identified because of Concerti’s unique integration of time-scaled trees.

## 4 Conclusion

In this paper we introduce Concerti, an algorithm for inferring evolutionary phylogenies. Concerti’s ability to extract and integrate information from multi-point, whether multi-site, multi-time, or combination thereof samples, enables the discovery of clinically plausible phylogenetic trees that capture the heterogeneity known to exist both spatially and temporally. These models can have direct therapeutic implications since they can highlight: “births” of clones that may harbor resistance mechanisms to treatment, “death” of subclones with drug targets, and acquisition on functionally pertinent mutations in clones that may have seemed clinically irrelevant. We demonstrate in this paper how Concerti can be applied to any genomic sequencing dataset with varying allele frequencies, whether it be cancer or the new SAR-Cov-2 virus causing the Covid-19 pandemic, and the results can have profound disease-specific clinical implication.

Identifying the presence of multiple viral strain infecting a single host can have significant impact on how we approach treatment, vaccine development, and mitigation strategies. The results for Covid-19 patients demonstrate Concerti’s ability to distinguish between viral strains based on difference allele frequencies and discover the presence of new homoplasies. Thus, Concerti’s results addresses the overwhelming challenges researches face when developing therapeutics and may help facilitate the key to effective vaccine development. Accurately monitoring tumor evolution over the course of a disease can lead to the identification of new drug targets and therapeutic approaches that can stabilize this complex disease and manage the selective pressures introduced by treatment exposure and tumor-environment changes. These results for patients CLL1, CLL2 and GI1 demonstrate how Concerti’s specific integration of multi-point data can facilitate better treatment plans that can both be more locally targeted and optimized for treatment responsivity.

## Data and Tool Availability

All data used in the paper are available with the original publications. In particular, patient CLL1 and CLL2 correspond to B06 and A43 of [13], respectively. While the GI1 colon cancer patient used for the multi-site analysis was previously published as TPS037 in [22]. The COVID-19 patient data come from 5 NCBI BioProjects: PRJNA601736, PRJNA603194, PRJNA610428, PRJNA605983 and PRJNA608651. The reads were trimmed with Trimmomatic (v. 0.39) and then mapped with bwa on MN908947.3 GenBank sequence. This sequence was taken in December from a 57 year old woman, who sold shrimp at the Wuhuan seafood market, appears to be the earliest case with COVID-19. Variant calling was performed generating mpileup files using SAMtools and then running VarScan (min-var-freq parameter set to 0.01). Finally to remove possibly sequencing artifacts, we retains SNV that: showing a VarScan significance p-value < 0.05 (Fisher’s Exact Test on the read counts supporting reference and variant alleles) and VAF>10%, resulting in a list of 41 SNVs in 21 patients that are used in the paper. Concerti’s binaries are available upon request.

## Acknowledgements

We are very grateful to Ignaty Leshchiner, Gad Getz, and Catherine J. Wu for providing CLL patient data.

